# VariantFlux: A genotype-first modelling workflow for predicting the impact of genetic variations on human metabolism

**DOI:** 10.64898/2026.08.03.740213

**Authors:** Hadi Nazem-Bokaee

## Abstract

Human genetic variation is a major determinant of organ metabolism, yet how naturally occurring variants shape quantitative metabolic phenotypes remains unclear. We present VariantFlux, a workflow that integrates ancestry-aware variant interpretation into genome-scale metabolic modelling to generate personalised, variant-constrained kidney reconstructions. Using the Human1 v1.19 model, we built a kidney-specific baseline model constrained by 482 metabolites and analysed 2,547 individuals from the 1000 Genomes Project, in whom ∼50% of metabolic genes were predicted damaging by at least three computational tools. These variants, whose burden differed subtly across ancestries, were translated into gene-dosage–anchored flux constraints for homozygous knockouts and graded heterozygous knockdowns. Despite widespread perturbation, >97% of models preserved baseline growth, indicating strong metabolic robustness. Yet individual genomes exhibited distinct flux-rewiring patterns, with frequent individual-specific gain-of-flux events and fewer shared loss-of-flux reactions. Limited ancestry clustering suggests metabolic responses are driven mainly by unique variant combinations. VariantFlux links human genomes to organ-level flux phenotypes, enabling precision medicine, pharmacogenomics, and disease risk prediction.

**Conceptual advance:** We present VariantFlux, a genotype-first framework that integrates predicted variant effects directly into genome-scale metabolic networks to generate personalised, organ-specific flux phenotypes. Unlike association-based metabolomics studies, this bottom-up approach enables exploratory, mechanistic prediction of how naturally occurring genetic variation reshapes human metabolism.

## Introduction

Despite remarkable progress in high-throughput and single-molecule genome sequencing^1,2^, multi-omics technologies^3^, and computational genomics^4,5^, a fundamental challenge persists in biomedical research: elucidating how upstream molecular perturbations—such as genetic variations—translate into downstream phenotypic consequences, including altered metabolic states and disease susceptibility. This complexity arises from the multilayered nature of gene regulation, the intricate interconnectivity of metabolic networks, cellular heterogeneity, and the current lack of unified frameworks capable of integrating genomic variant data into mechanistic biological predictions. Bridging these gaps is essential for understanding how genetic variation shapes molecular physiology, disease risk, and therapeutic response across individuals and populations.

Population-scale genome sequencing efforts, such as the UK Biobank study of over half a million participants, have revealed that each healthy individual carries approximately five million genetic variants, including numerous potentially deleterious ones^6^. On average, a person may harbor over 400 variants predicted to be damaging, with several carrying pathogenic potential during their lifetime^7^. Moreover, healthy individuals can carry about 100 predicted loss-of-function (LoF) variants, including complete inactivation of roughly 20 genes, yet most remain asymptomatic^8^. This observation underscores a key gap in our understanding—why some variants exert strong phenotypic effects while others do not. Variant penetrance varies widely across genes and biological contexts^9^: haplo-insufficient genes are extremely sensitive to dosage reduction, while others, such as olfactory receptor genes, tolerate complete loss of function. Natural selection tends to purge highly deleterious alleles, rendering truly pathogenic variants extremely rare. As a result, elucidating the metabolic and physiological consequences of rare damaging variants demands genotype-first approaches applied to very large, functionally interpretable cohorts.

Another layer of complexity lies in the diversity of variant effect prediction tools, which rely on distinct principles and sometimes yield inconsistent predictions^10^. Furthermore, most genome-wide association studies (GWAS) have been conducted in European populations, introducing ancestry biases into variant catalogues and risk prediction models^11^. This limitation is particularly acute for underrepresented populations^12^, where population-specific variant databases and reference genomes remain incomplete. Even as population-specific reference genomes advance variant cataloguing, systematic approaches will still be required to assess functional impact of variants in a mechanistic, context-dependent manner.

Finally, many complex diseases cannot be explained by alterations in single genes alone but instead arise from the combined, often subtle effects of many variants across multiple loci^13,14^. Studies employing multi-ancestry polygenic risk scores have shown that the cumulative influence of hundreds of genetic loci can drive susceptibility to diseases such as kidney and type 2 diabetes across diverse populations^14^. This polygenic, multi-target nature of diseases highlights the need for frameworks that can model the distributed and nonlinear effects of multiple variants acting simultaneously.

Recent research has increasingly focused on intermediate molecular phenotypes that bridge genetic variation and disease, with metabolites emerging as particularly informative readouts of gene function^15–17^. Metabolite levels are strongly influenced by genetic variation near metabolic enzymes and transporters, and metabolome-wide association studies (mGWAS) have uncovered numerous gene–metabolite links across human populations^18–20^. Notably, recent large-scale efforts combining exome sequencing with population metabolomics have demonstrated that even rare damaging heterozygous variants can exert graded, measurable effects on metabolite concentrations, challenging the traditional view of heterozygosity as functionally silent^19^. These studies provide compelling empirical evidence that naturally occurring variants shape human metabolism in subtle but quantifiable ways.

Despite these advances, association-based approaches remain fundamentally constrained by the availability of matched genomic and metabolomic data and by the metabolites that can be experimentally measured. Moreover, metabolite concentrations represent static snapshots of cellular state and do not directly capture metabolic activity, which is governed by flux—the rates of biochemical transformation and transport through metabolic pathways. As a result, many genotype–metabolite associations remain difficult to interpret mechanistically, and causal links between genetic variation and metabolic function are often inferred rather than predicted.

Genome-scale metabolic models (GSMMs) offer a complementary, mechanistic framework to address these limitations. By explicitly encoding the stoichiometry, connectivity, and gene–reaction relationships of metabolic networks, GSMMs enable prediction of flux distributions under genetic or environmental perturbations^21,22^. Importantly, they can integrate the combined effects of multiple variants across pathways, capturing emergent, systems-level behaviour that cannot be inferred from single-gene associations alone^19,23^. While GSMMs have been widely applied to study disease states and tissue-specific metabolism, they have rarely been used in a systematic, genotype-first manner to explore how naturally occurring human variation reshapes metabolic function across healthy individuals.

Here, we introduce Variant-informed metabolic Flux analysis (VariantFlux), a bottom-up framework that mechanistically links potentially damaging genetic variants to metabolic network behaviour. VariantFlux integrates computational variant-effect predictions directly into a GSMM by imposing gene-dosage constraints, generating personalised, variant-constrained metabolic reconstructions without reliance on prior metabolomic measurements or known genotype–phenotype associations. This approach enables systematic identification of dosage-sensitive and recessive metabolic genes, characterisation of network-level flux rewiring, and prioritisation of functionally relevant metabolic vulnerabilities (**Figure 1**).

**Figure 1.**
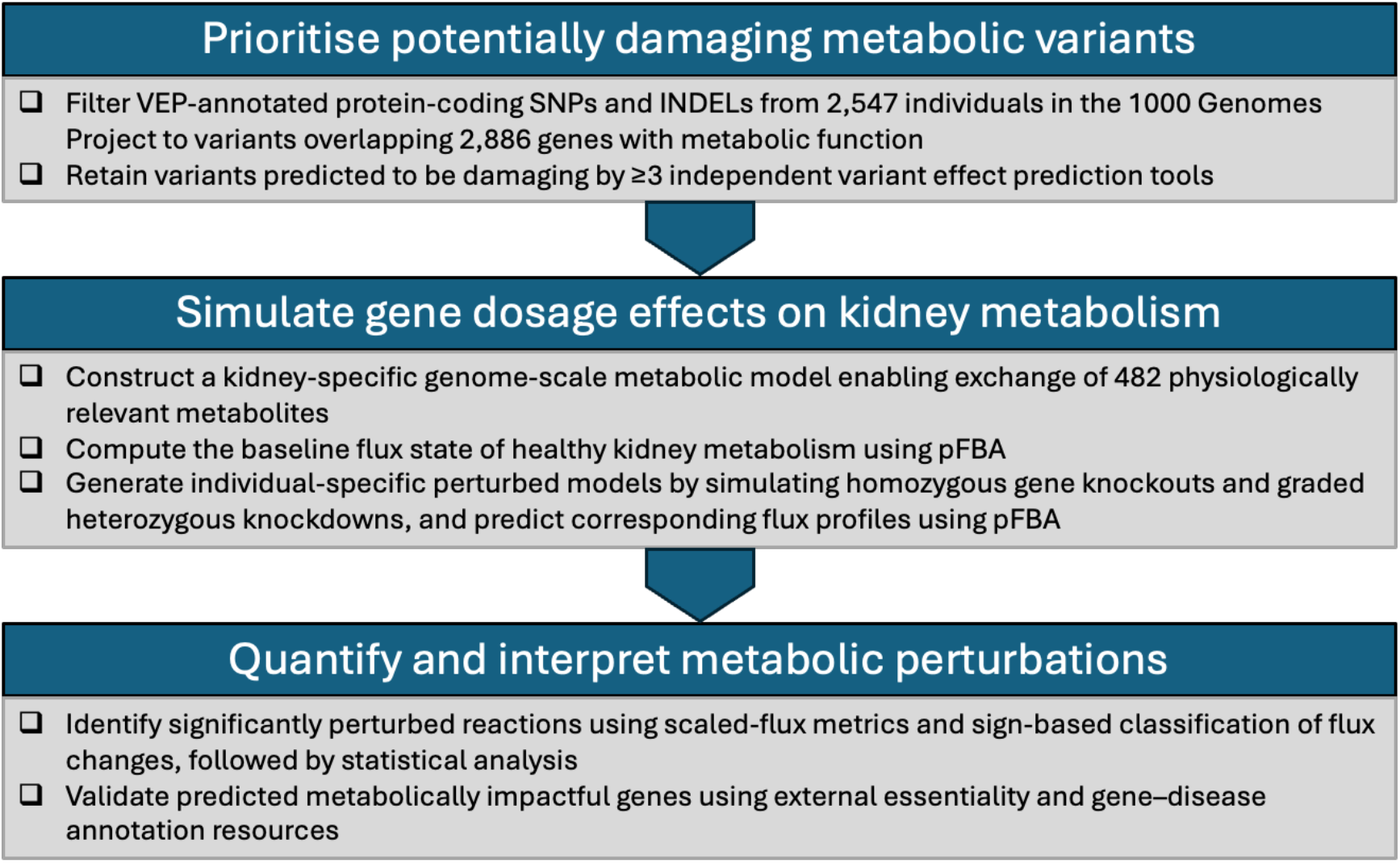
VariantFlux proposed workflow. The VariantFlux framework enables downstream applications including prioritisation of candidate metabolic genes for biomarker development and therapeutic targeting, as well as iterative model refinement through experimental validation.

To demonstrate the utility of VariantFlux in linking genetic variation to organ-specific metabolism, we applied the framework to human kidney metabolism. We constructed a kidney-specific GSMM derived from the Human1 v1.19 reconstruction, constrained using physiologically relevant urinary and circulating metabolites to represent a healthy renal baseline. Predicted damaging variants from 2,547 individuals in the 1000 Genomes Project were then mapped onto this network as gene-specific dosage perturbations, enabling systematic simulation of homozygous knockouts and graded heterozygous knockdowns and the generation of individual-specific, variant-constrained kidney metabolic models.

Using flux balance analysis, we quantified the impact of genetic perturbations on growth capacity, reaction-level fluxes, and pathway usage, revealing extensive yet structured metabolic rewiring across individuals. This genotype-first framework captures both shared network constraints and highly individual-specific responses, providing a scalable foundation for predictive, ancestry-aware metabolic modelling. While demonstrated here in the kidney, VariantFlux is readily transferable to other tissues or populations, enabling hypothesis generation in disease biology, biomarker discovery, and pharmacogenomics beyond association-based inference.

## Methods

### Data collection and pre-processing

Single-nucleotide variants (SNVs) and insertions/deletions (INDELs) relative to the GRCh38 reference genome for 2,547 individuals from the 1000 Genomes Project were obtained from the International Genome Sample Resource (IGSR) web source^24^. Variants overlapping protein coding (CDS) regions––based on Ensembl annotations (v.114)^25^––were extracted using bcftools view^26^, producing a reference set of all protein-coding genes.

### Variant effect prediction using the Ensembl Variant Effect Predictor

We annotated coding variants using Ensembl VEP ^27^ (v114, vep -sift b -polyphen b - plugin pLI, AlphaMissense). Transcript-level effects were predicted with SIFT^28^ and PolyPhen^29^; gene-level loss-of-function intolerance with pLI^30^ plugin; and missense pathogenicity with AlphaMissense^31^. Intergenic and intronic variants were excluded from the analysis.

### Variant prioritisation in metabolic genes

To focus on genes with metabolic function, we extracted 2,886 metabolic genes from the Human1 genome-scale metabolic model reconstruction^32^ (v1.19.0) and mapped their Ensembl IDs to Ensembl v114 GFF3 annotations to obtain genomic coordinates. This reference table was used to filter VEP-annotated variants to those overlapping the target metabolic gene set.

Damaging variants were then prioritised using a custom Python workflow. Variants were retained if they overlapped a metabolic gene and were heterozygous or homozygous for alternate alleles. A variant was classified as damaging if it met at least three of the following cut-off criteria: SIFT = deleterious, PolyPhen = probably damaging, AlphaMissense score > 0.564, pLI > 0.9, or ClinVar = pathogenic.

Because genomic variants can occur in regions that influence more than one gene, some variants may be non-damaging for a metabolic gene in our set while still affecting a nearby or overlapping non-metabolic gene. Such cases were noted but not prioritised, as the analysis specifically targets functional impacts within the curated metabolic gene list.

Individual-level summaries of damaging variants in metabolic genes were generated (**Supplementary Table S1**), and these profiles were used to inform subsequent simulation and knockout analyses in the kidney-specific metabolic model.

### Assembly of the baseline kidney-specific metabolic model

We constructed a kidney-specific baseline model (*Human1BL*) using the Human1 v1.19 genome-scale metabolic reconstruction (8,363 metabolites, 12,971 reactions, and 2,887 genes across 139 pathways). The generic Human1 metabolic model (*Human1G*) served as the starting point. To tailor the model to kidney physiology, we first identified the set of exchange metabolites relevant to renal function.

Urinary metabolites were identified by parsing Human Metabolome Database (HMDB) v5.0^33^ XML files and selecting metabolites annotated as “urine” in the *biofluid*, *biospecimen*, *normal_concentrations*, or *abnormal_concentrations* fields. Identified metabolites were then mapped to the *Human1G* reconstruction using multiple identifier systems, including ChEBI, KEGG, PubChem, MetaNetX, LipidMaps, EHMN, and HMR2. A curated list of circulating metabolites in proximal tubule epithelial cells was compiled from prior studies^17,34^ and cross-checked against metabolites in the renal objective function of Chang et al^35^ and the HEK cell model of Quek et al^36^. Metabolites lacking an extracellular representation in *Human1G* were excluded. After confirming the final set of kidney-relevant exchange metabolites, bounds of all other exchange reactions were fixed to zero, and the generic *Human1G* biomass reaction was retained to represent renal cellular composition^37,38^. The resulting *Human1BL* model (**Supplementary File 1**) was used for all simulations of kidney metabolism under healthy or perturbed conditions as described below.

### Simulation of heterozygous and homozygous loss of gene function at individual-level

We systematically assessed the impact of genetic variants on kidney metabolism by performing single-gene knockout (homozygous state) and knockdown (heterozygous state) simulations to generate individual-specific metabolic flux profiles. Essential genes were identified through in silico single-gene knockouts using the *Human1G* Gene-Protein-Reaction (GPR) rules, where loss of any required subunit disables the associated reaction^39^. Genes whose deletion abolished biomass production were designated as essential, accounting for isozyme redundancy. Reactions linked to these essential genes were retained in all subsequent simulations, ensuring model feasibility by preventing their corresponding genes from being targeted for knockout.

For each individual, homozygous variants were simulated as full knockouts by constraining all associated reactions to zero flux. Heterozygous variants were modelled as knockdowns by scaling reaction bounds to 50% fraction of baseline flux while preserving flux directionality. Objective-reverse reactions were blocked to prevent artefactual flux redistribution.

### Identification and classification of perturbed ffuxes

Flux deviations of each impacted reaction were quantified using a bounded, symmetric scaled-flux distance,

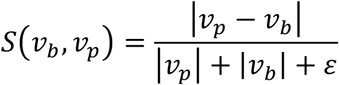

where *v_b_* and *v_p_* are baseline and perturbed fluxes, respectively, and *ε*=10^−9^ prevents division by zero. This metric, which ranges from 0 to 1, identifies reactions with *S*>0.5 as significantly perturbed. The formulation is conceptually related to Canberra distance^40–42^––with slight modifications––which normalises pairwise differences by the sum of magnitudes and is well suited for data with zeros, large dynamic ranges, and asymmetric distributions—properties characteristic of metabolic fluxes. Normalisation ensures that reactions with large absolute fluxes do not dominate, biologically meaningful changes in low-flux reactions are preserved, and flux differences are expressed on an interpretable scale.

Significantly perturbed reactions were further assigned to one of four flux-type categories based on sign and activity patterns: Complete Loss (CL) for *v_b_* ≠ 0 and *v_p_*= 0; Complete Gain (CG) for *v_p_* ≠ 0 and *v_b_*= 0; Flux Switch (FS) for *v_b_ • v_p_* < 0, indicating reversal of direction; and Partial Change (PC) for all other cases involving non-zero fluxes of the same sign but altered magnitude. Together, the scaled-flux metric and sign-based rules provide a robust and interpretable framework for detecting and classifying diverse types of flux perturbations.

Flux distributions were computed using parsimonious Flux Balance Analysis (pFBA)^43^, with reactions exhibiting significant changes further interrogated using Flux Variability Analysis^39^ (FVA). All models (*Human1G*, *Human1BL*, and individual-specific kidney models) were expressed in FBA-compatible units^44^ (mmol/gDCW·h; biomass h⁻¹) and fluxes were normalised by biomass to remove growth-rate bias. Simulations were implemented using COBRApy toolbox^45^ with the GLPK solver, and parallelised execution enabled efficient processing of all 2,547 samples. Outputs included sample-specific flux distributions, reactions with significant changes, and optional synthetic lethal predictions.

### Statistical analyses and reproducible workffow

Differences in the distribution of predicted damaging gene counts and relative biomass flux across ancestry groups were assessed using the Kruskal–Wallis test, selected for its robustness to non-normal distributions and unequal group sizes. Group-wise distributions were visually inspected using violin plots to confirm comparable spread and skewness prior to hypothesis testing. To further quantify ancestry effects on predicted damaging gene burden and biomass growth, generalised linear models (GLMs) with a Poisson family and log link were fitted, with ancestry included as a categorical predictor.

All statistical analyses and figure generation were performed in Python using SciPy for non-parametric testing and Statsmodels for GLM fitting. Analyses were implemented within a modular, configuration-driven computational workflow executed in Jupyter, in which dataset-specific inputs and file paths were defined externally via a YAML configuration file. This design enabled systematic analysis of multiple datasets using a single, shared codebase and allowed all statistical results and figures to be deterministically regenerated by re-executing the notebook with the corresponding configuration file.

## Results

### 1. Distribution of damaging metabolic variants across ancestry groups

A total of 11,303 unique CDS regions harboured one or more predicted damaging variants (**Figure 2A**). Of the 2,887 metabolic genes analysed, three autosomal genes (*ATP5F1EP2*, *SLC37A4*, and *USP41*) and 13 mitochondrial genes showed no overlap with annotated CDS regions and were excluded. Among the remaining 2,871 genes, 1,848 (64.3%) contained at least one damaging variant (**Figure 2B**), and 1,426 (49.6%) were predicted as damaging by SIFT, PolyPhen, and AlphaMissense (**Supplementary Figure S1**).

**Figure 2.**
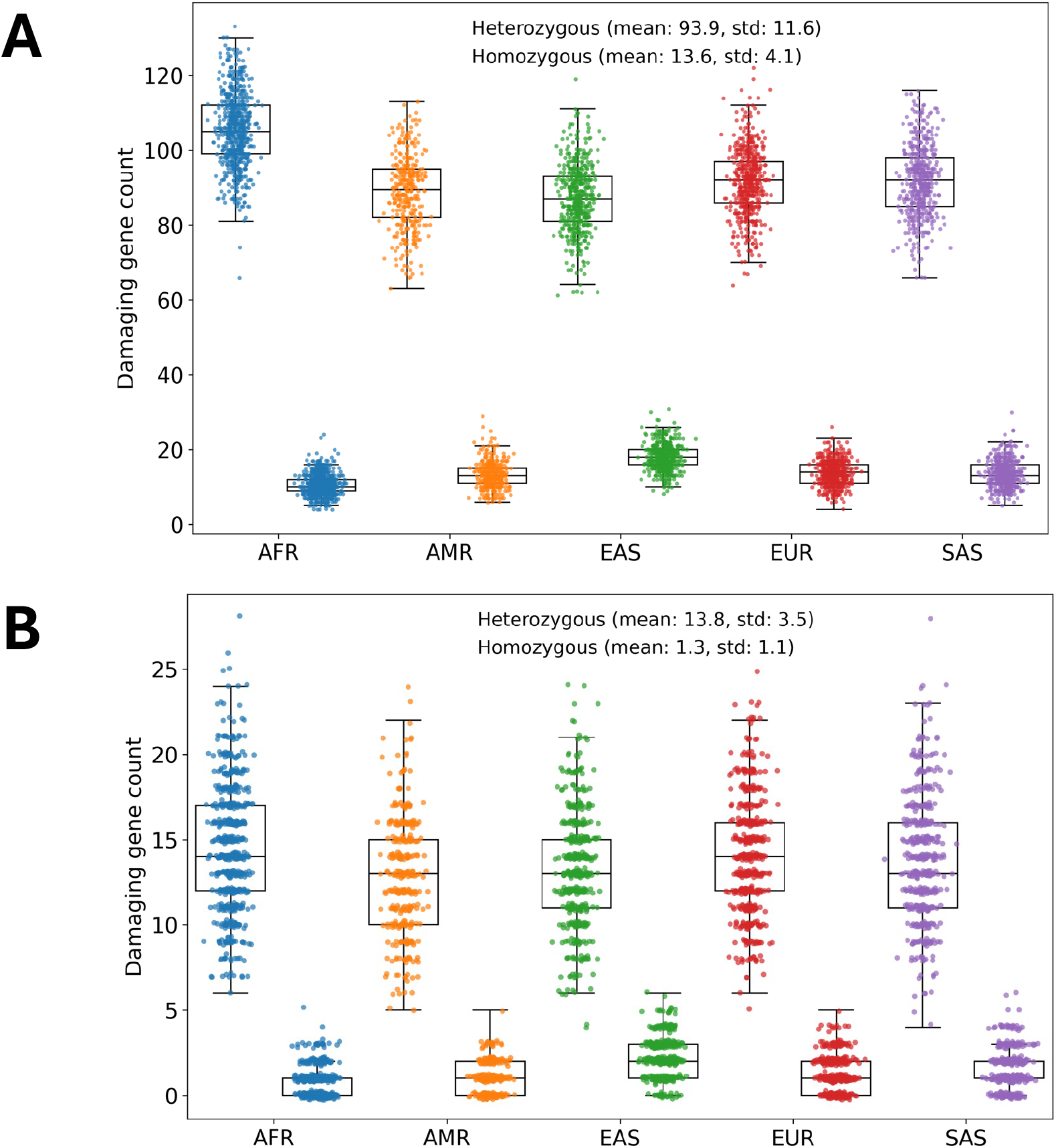
Distribution of genes predicted to carry damaging variants across ancestry groups (N = 2,547 individuals). **Panel A** shows the results for 11,303 protein-coding human genes, and **Panel B** shows results for genes with known metabolic functions from the Human-GEM model (2,886 total genes). Heterozygous and homozygous states (and their corresponding mean and standard deviation of gene counts) are shown by top and bottom box plots, respectively. Each dot represent a single individual for which only genes predicted as damaging by at least three independent variant effect prediction tools across all samples are shown. AFR: African ancestry (671 individuals); AMR: American ancestry (348 individuals); EAS: East Asian ancestry (515 individuals); EUR: European ancestry (521 individuals); SAS: South Asian ancestry (492 individuals).

Across individuals, the mean number of metabolic genes carrying at least one damaging variant was approximately 15 per genome (**Figure 2**). A Kruskal–Wallis test across the five continental ancestry groups revealed that at least on group differed in overall damaging gene counts (H = 45.47, p = 3.17 x 10^-9^). Bootstrap resampling of individuals (1,000 iterations, subsample size = 300 per ancestry group) confirmed that the observed ancestry differences in damaging gene counts were robust, with median p-values of 9.77 x 10^-9^ and 3.18 x 10^-65^, for heterozygous and homozygous variants, respectively, and 100% of iterations remaining significant. (**Supplementary Table S2**).

Focusing on heterozygous variants, individuals of African ancestry (AFR) had the highest median number of damaging genes (∼14.5 per genome). Relative to the AFR group, a generalized linear model (Poisson distribution) indicated that individuals from other ancestries carried slightly fewer damaging genes: AMR (∼12% lower), EAS (∼9%), SAS (∼5%), and EUR (∼4%). These trends were statistically significant (p < 0.01, **Supplementary Table S3-4**), consistent with observed Kruskal–Wallis differences.

### 2. Metabolic robustness under gene dosage perturbation predicted by growth rate

Parsing HMDB yielded 1,618 urinary metabolites, which collapsed to 467 unique extracellular metabolites in *Human1G*. The curated list of 93 circulating proximal tubule metabolites overlapped substantially with the urinary set, with 78 shared and 15 unique. After cross-referencing with previous renal models, these contributed to a final set of 482 kidney metabolites.

Using the *Human1BL* baseline kidney model, we simulated combined homozygous knockouts and heterozygous knockdowns for all 2,547 individuals. Approximately 33% of perturbed kidney models (847 individuals) retained baseline growth rates (i.e., biomass fluxes), indicating metabolic resilience to partial gene loss (**Figure 3A**). The remaining ∼66% kidney models exhibited reduced growth, suggesting dosage-sensitive genes whose partial inactivation impairs metabolic capacity. Generalized linear modelling revealed that relative growth rates were largely comparable across ancestry groups, with only South Asian individuals showing a modest but statistically significant reduction compared to the African reference group (p = 0.049).

**Figure 3.**
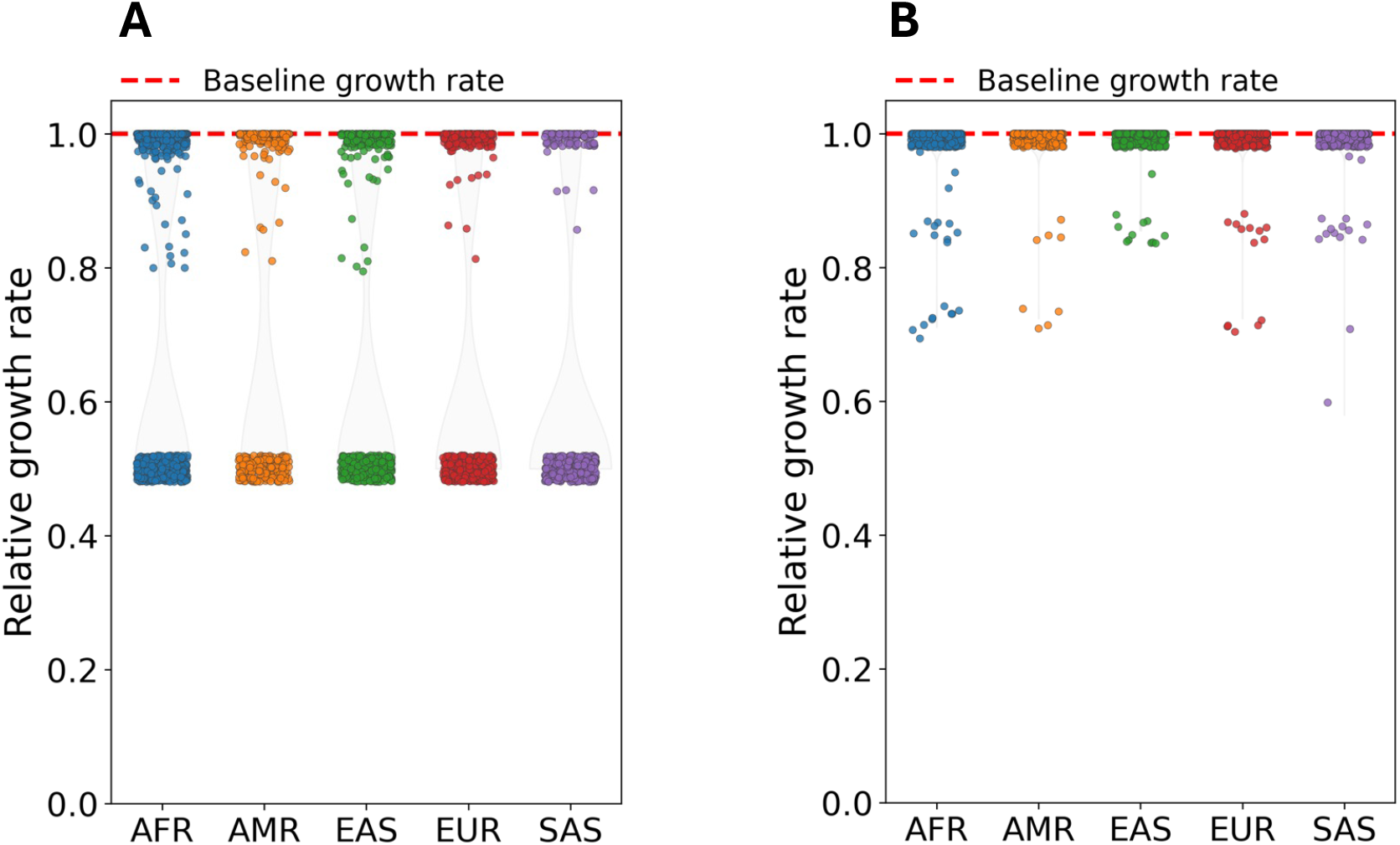
Predicted relative growth rate (i.e., biomass flux) of perturbed kidney models for 2,547 individuals across ancestry groups. Relative biomass fluxes were calculated by dividing the biomass flux predicted by the perturbed model to that of the unperturbed baseline model––shown as red dashed lines––for each individual. Each dot represents one individual. Each violin plot shows the combined effect of full gene knockout for homozygous state and 50% knockdown (**Panel A**), or full knockout (**Panel B**) for heterozygous state for the individuals in the respected ancestry group.

When simulating gene knockouts across both heterozygous and homozygous states, no individual-specific kidney model from any ancestry group exhibited a complete loss of growth. Under full gene knockouts, at least 97% of the 2,547 perturbed kidney models retained baseline-level biomass flux, while the remaining few showed reductions in growth (**Figure 3B**). Generalized linear modelling of relative biomass flux across ancestry groups revealed no significant ancestry-dependent differences (p > 0.95 for all comparisons; pseudo-R² = 8.8 x 10^-7^), indicating that predicted kidney model growth rates were consistent across populations. Analysis of the <3% of individuals showing reductions in relative growth rate at the sub-population level indicated that such decreases were rare across ancestry groups, affecting 1.34% of individuals in AFR, 1.15% in AMR, 0.95% in EUR, and 0.4% in SAS populations. Given their small effect size and limited frequency, no further investigation into the underlying causes was pursued. (**Supplementary Figure S2**).

### 3 Network-level variant impact beyond biomass

In the *Human1BL* model, 630 reactions (∼5%) carried non-zero flux based on pFBA, and 2,894 (∼29%) were flux-flexible predicted by FVA. Following knockout of homozygous variants and knockdown of heterozygous variants, 1,771 reactions exhibited significant perturbation (scaled flux > 0.5), while combined knockout of both variant types affected 1,139 reactions.

Principal component analysis (PCA) of flux-type proportions for significantly perturbed reactions revealed a primary axis of variation capturing large-scale network reorganisation beyond growth effects (**Figure 4A**). Across the perturbed kidney models following knockout of homozygous variants and knockdown of heterozygous variants, the first two components explained 56.4% of the total variance, with PC1 (31.6%) primarily separating impacted reactions dominated by complete gain (CG) versus complete loss (CL) flux-type behaviours. CL showed strong positive loadings and CG negative loadings on PC1, highlighting a major gain–loss contrast in reaction-level flux patterns. Consistent with these loadings, CG-dominant reactions were numerous (n = 1,145) but observed in relatively few individuals (median = 5 individuals), whereas CL-dominant reactions were fewer (n = 252) yet widespread (median = 1,689 individuals). Reactions classified as partial change (PC; n = 215) or flux switch (FS; n = 159) exhibited intermediate or localised patterns (median = 1,652 and 4 individuals, respectively). Shannon entropy values mirrored these trends, with CL (0.26) and PC (0.25) showing greater variability across individuals compared to CG (0.00003) and FS (0.03).

**Figure 4.**
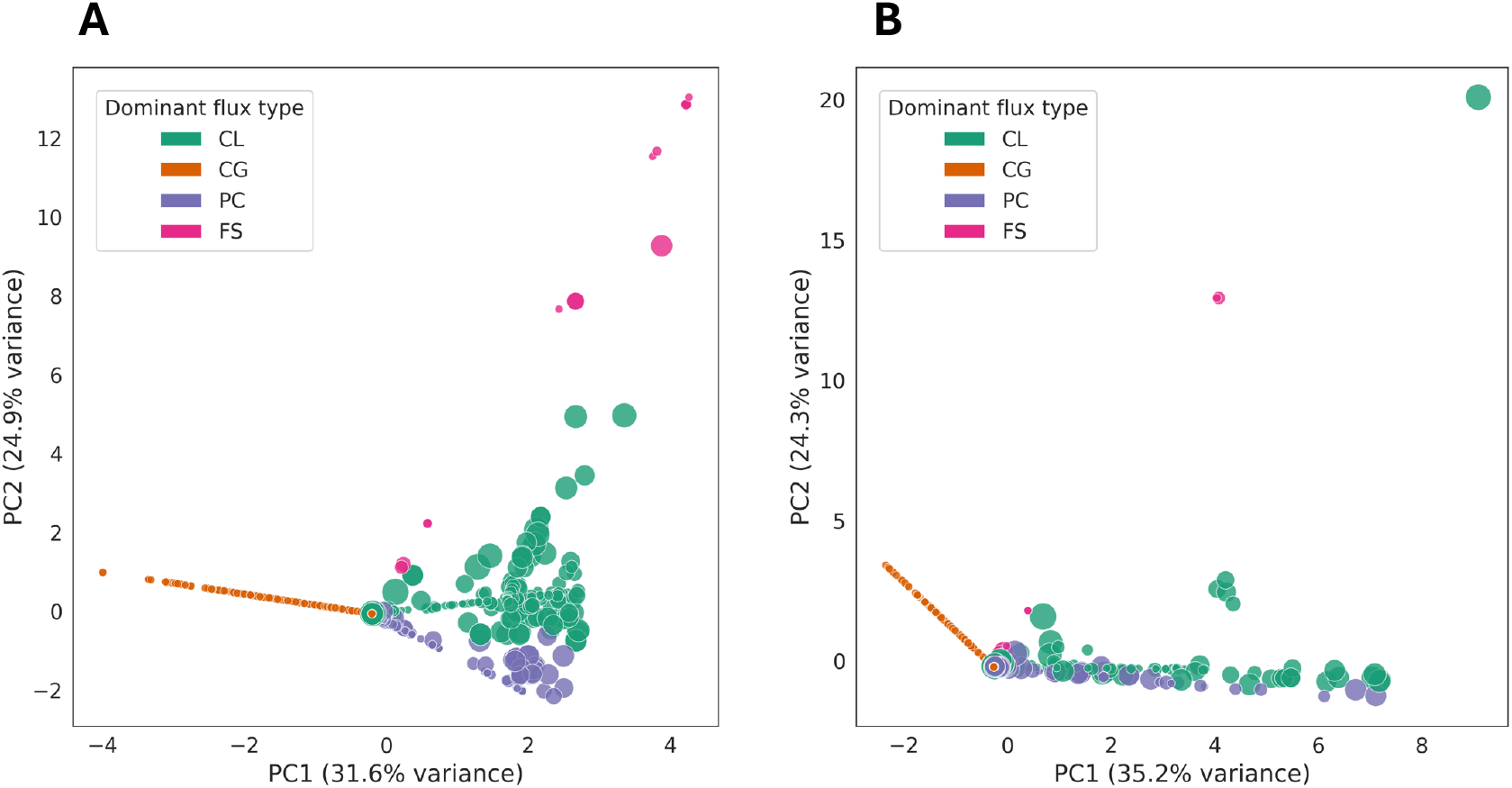
Principal Component Analysis (PCA) of reaction-level flux perturbations across all kidney metabolic models. **A.** PCA of flux-type proportions for all reactions with significant flux perturbation (scaled flux > 0.5) following knockout of homozygous variants and knockdown of heterozygous variants. PC1 (31.6% variance) separates reactions dominated by complete gain (CG) versus complete loss (CL). CG-dominant reactions are numerous but sample-specific, whereas CL-dominant reactions are fewer and widespread. Partial change (PC) and flux switch (FS) reactions show intermediate patterns. Dot size represents Shannon entropy, indicating the heterogeneity of flux-change types per reaction (larger dots = more heterogeneous). **B.** PCA following combined knockout of homozygous and heterozygous variants shows a similar structure. CG-dominant reactions remain largely sample-specific, CL-dominant reactions are more prevalent, and PC/FS reactions exhibit higher heterogeneity. These patterns indicate that population-level metabolic perturbations are primarily driven by loss- or reduction-type flux changes, while gain-type changes tend to be localised or compensatory. Dot size again reflects entropy per reaction.

Analysis of reactions impacted by combined gene knockout of homozygous and heterozygous variants revealed a similar structure (**Figure 4B**): CG-dominant reactions (n = 675) occurred in relatively few individuals (median = 16), whereas CL-dominant reactions (n = 263) were fewer but more widespread (median = 49). PC (n = 182) and FS (n = 19) reactions were intermediate in prevalence and exhibited higher entropy, consistent with more heterogeneous flux responses.

In addition to the global analysis of significantly perturbed reactions, we specifically examined reactions directly associated with the perturbed genes, regardless of whether they exceeded the scaled-flux threshold for significance. Among individuals whose kidney models showed ≥5% reduction in biomass flux compared to baseline growth, 1,112 individuals (43.6%) in the homozygous knockout plus heterozygous knockdown simulations had at least one directly associated reaction affected (ancestry distributions: AFR 23.9%, AMR 12.2%, EAS 19.9%, EUR 21.4%, and SAS 22.6%). In the combined knockout analysis, 67 individuals (2.6%) were affected, with a similar ancestry distribution pattern (AFR 29.9%, AMR 11.9%, EAS 17.9%, EUR 20.9%, and SAS 19.4%) (**Figure 5A**).

**Figure 5.**
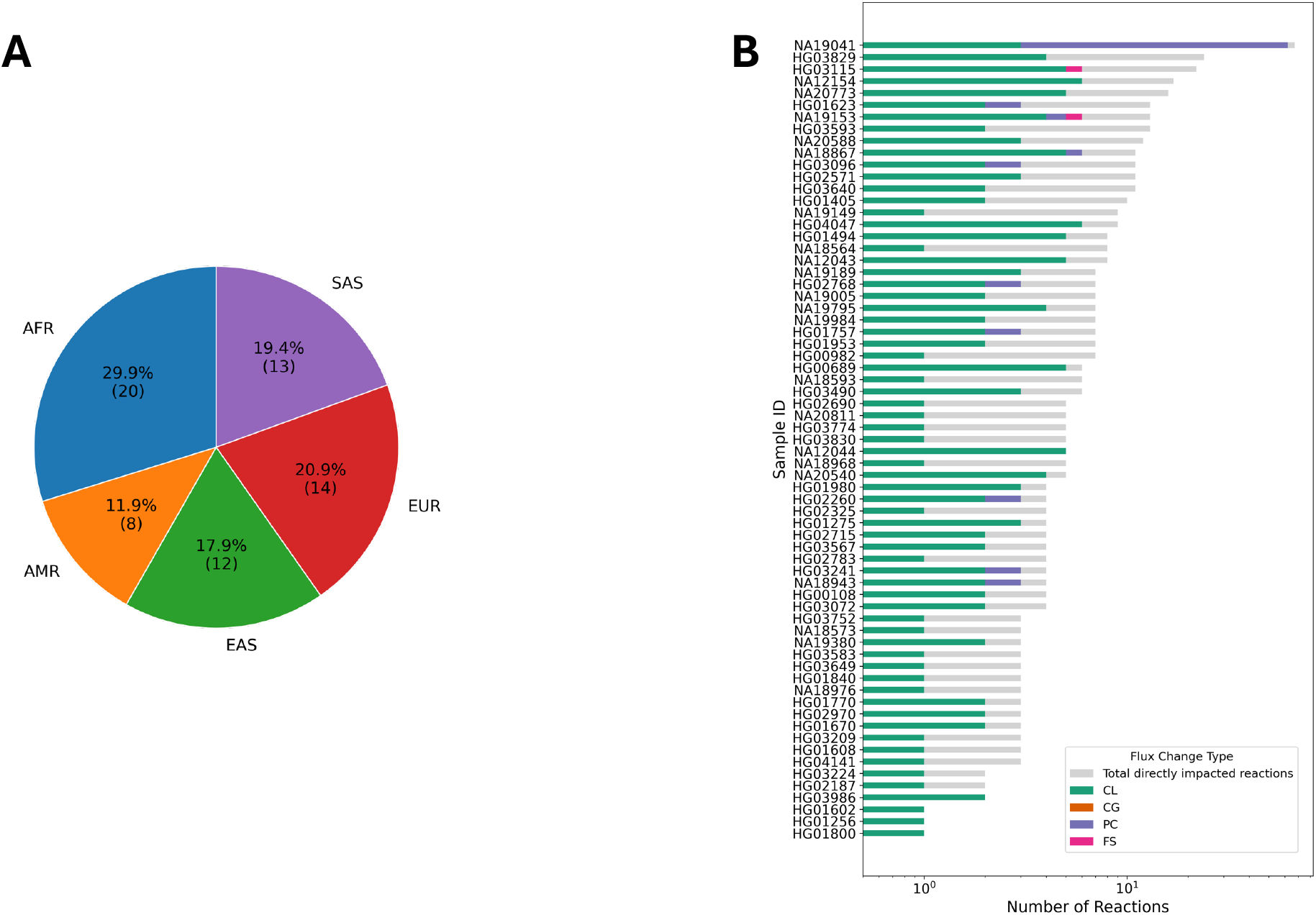
Distribution of reactions directly impacted by gene perturbations for individuals with ≥5% reduction in the predicted relative biomass flux across ancestry groups. (**A**) Pie chart showing the distribution of individuals (total of 67) across ancestry groups (AFR, AMR, EAS, EUR, SAS) for the combined homozygous and heterozygous knockouts simulations. (**B**) Stacked bar chart showing the number of reactions directly affected by gene knockouts per individual (combined homozygous and heterozygous knockouts), grouped by flux-type change: complete loss (CL, green), complete gain (CG, orange), partial change (PC, purple), flux switch (FS, pink), and no significant change (NA, gray). Samples are ordered by total number of directly impacted reactions.

When aggregated across unique reactions, the distribution of flux-type changes differed between simulation sets. In the homozygous knockout plus heterozygous knockdown simulations, CL was the most frequent category (159 reactions), followed by PC (n=115), and FS (n=10). By contrast, in the combined knockout simulations, PC predominated (65 reactions), with CL (n=36), and FS (n=1) reflecting smaller perturbations overall. The flux through the remaining gene-associated reactions remained unchanged despite perturbation (n=136 and 64 for homozygous knockout plus heterozygous knockdown simulations and combined knockout simulations, respectively). The number of impacted gene-associated reactions per individual ranged from a single reaction to more than 60, reflecting notable variability in metabolic consequences across individuals. For example, HG01800, HG01256, and HG01602 displayed predominantly CL events, whereas NA19041 exhibited predominantly PC events (n = 59) (**Figure 5B**).

### 4. Identification of clinically relevant candidate genes predicted to impact kidney metabolism

To assess whether model-predicted metabolic perturbations correspond to known biological or clinical relevance, we compared essential genes identified by single-gene knockout simulations to experimental essentiality databases. Of the 105 genes predicted to be essential for growth in the kidney-specific model, 44 (42%) were annotated as essential in the Online Gene Essentiality Database^46^ (OGEE). Additional comparisons with other gene-disease resources further supported several model predictions, including those in core energy, amino acid, and transport pathways. Searching 128 genes––that were predicted to be involved in full reaction inactivation across individuals for both full and partial loss of function conditions––in the Gene Curation Coalition^47^ and PanelApp^48^ resources, using the Human Phenotype Ontology^49^ terminology resulted in identification of 64 genes with kidney-related evidence (**Figure 6**).

**Figure 6.**
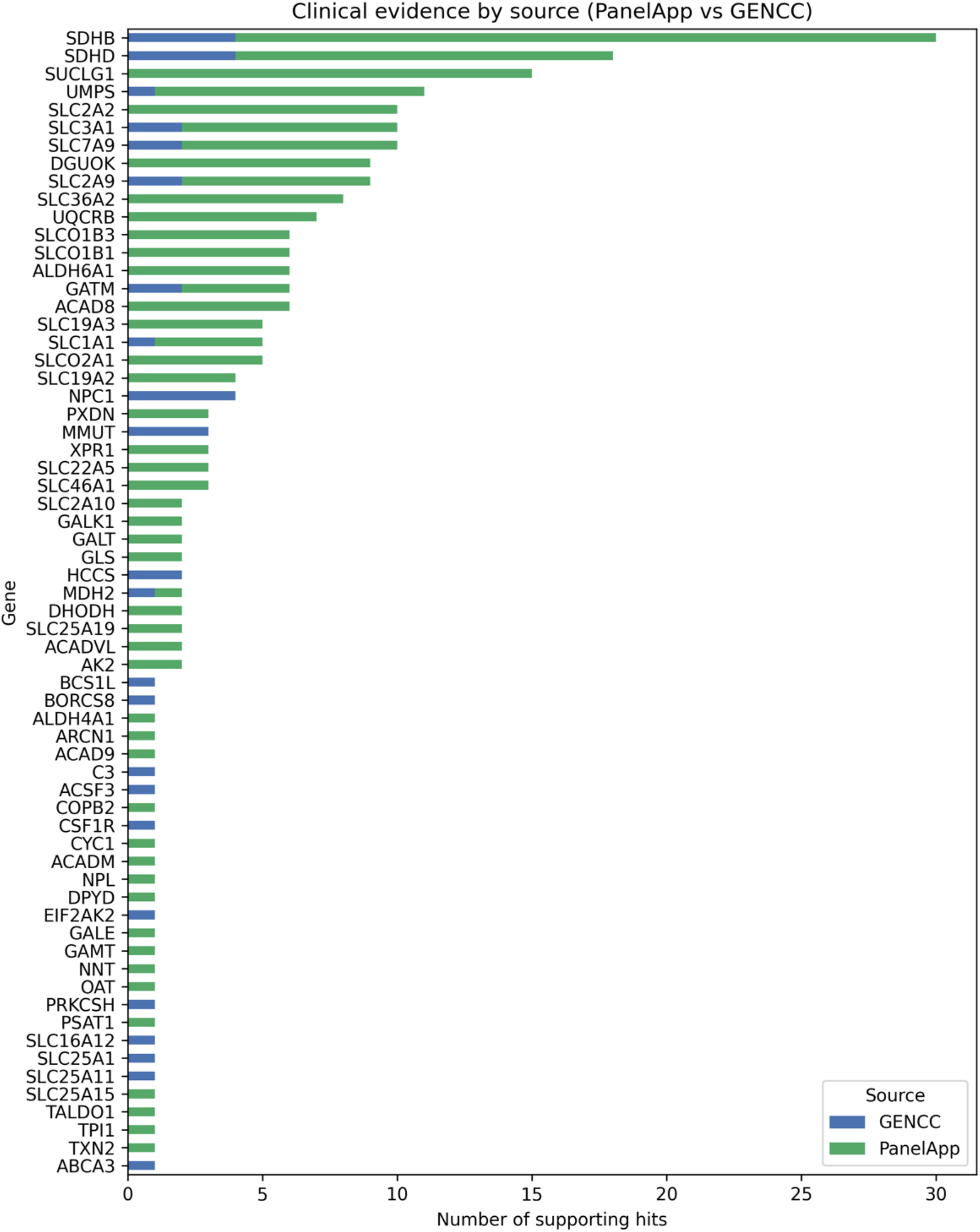
Stacked bar plot showing clinical evidence supporting metabolic genes predicted by the kidney model to undergo reaction inactivation across individuals under both full and partial loss-of-function conditions. The number of supporting hits indicates how many times each predicted damaging gene was found in either the GenCC database (20,033 gene entries) or the PanelApp resource (439 clinical panels) with phenotypic annotations matching any of 1,018 kidney-related Human Phenotype Ontology (HPO) terms.

## Discussion

The VariantFlux workflow integrates ancestry-aware variant interpretation directly into a genome-scale metabolic model, using damaging variant predictions to impose gene dosage constraints and thereby generate personalised, variant-constrained metabolic reconstructions. The results demonstrate that although damaging variants in metabolic genes are widespread across individuals and ancestries, kidney metabolism exhibits extensive buffering capacity that limits their impact on global metabolic performance. At the same time, VariantFlux exposes substantial, individual-specific flux rewiring that remains undetectable from growth phenotypes alone, highlighting the value of mechanistic modelling for disentangling genotype–phenotype relationships.

Across 2,871 metabolic genes analysed, approximately half carried variants predicted as damaging by at least three variant effect prediction tools (namely, SIFT, PolyPhen, and AlphaMissense). Although the overall shapes of variant-count distributions were similar across ancestry groups, statistical analyses revealed subtle yet robust ancestry-associated shifts in central tendency. Individuals of African ancestry carried slightly higher numbers of damaging heterozygous variants, in agreement with established patterns of greater genetic diversity^50,51^. Bootstrap resampling confirmed that these differences reflect genuine population-level distinctions rather than artefacts of sampling or distributional skew. These findings echo population-scale sequencing studies reporting that healthy individuals typically harbour dozens of genuine loss-of-function alleles, and underscore the need for mechanistic models (such as GSMMs) that incorporate population diversity rather than relying solely on a single reference genome. Without such adjustments, population-specific metabolic features relevant for disease susceptibility or pharmacogenomic response may be systematically overlooked.

Despite the prevalence of damaging variants, metabolic capacity of the kidney proved remarkably robust. More than 97% of individualised models maintained baseline biomass flux under both heterozygous and homozygous knockout conditions. This robustness, consistent across ancestry groups, likely reflects widespread pathway and isoenzyme redundancy within the human metabolic network and demonstrates that the kidneys’ essential functions—including solute transport, amino acid turnover, and mitochondrial ATP production—are buffered against single-gene perturbation^32^. Our genotype-informed knockdown strategy, which anchors heterozygous simulations to baseline fluxes assuming a linear relationship and prevents artefactual pathway reversals, further supports the inference that human kidney metabolism is buffered against modest reductions in enzyme capacity. This constraint-based method does not explicitly model higher-order regulatory compensation, instead it captures the network-level consequences of gene dosage changes. However, such buffering may mask latent vulnerabilities that only emerge under physiologically relevant stresses. Incorporating context-specific constraints such as hypoxia, substrate limitation, or disease-derived transcriptomic states may reveal stress-dependent sensitivities with clinical relevance. Beyond global growth effects, reaction-level perturbations revealed extensive metabolic reorganisation. Thousands of reactions exhibited significant flux changes, demonstrating that genetic variants propagate through the network in complex, nonlinear ways even when biomass output is maintained. Principal component analyses revealed a dominant gain–loss axis of variation, with complete loss (CL) and complete gain (CG) flux-types driving global metabolic differences across individuals. CL events were fewer but widespread across individuals, reflecting loss of flux through universally non-redundant reactions. In contrast, CG events were more numerous yet highly individual-specific, arising only in individuals whose remaining intact pathways support compensatory rerouting—highlighting how alternative metabolic routes become accessible in certain genetic backgrounds but not others, providing candidates for further analysis of gain-of-function mechanisms). Partial changes and flux-switches captured intermediate states of local rewiring and displayed higher entropy, indicative of heterogeneous responses across individuals. These patterns indicate that metabolic resilience is achieved through a mixture of ubiquitous and personalised compensatory mechanisms, producing distinct network-level signatures of gene dosage perturbation.

Analysis of reactions directly associated with perturbed genes further highlighted the personalised nature of metabolic responses. While only a small subset of individuals exhibited measurable reductions in biomass flux, a much larger fraction displayed direct gene-linked flux changes, sometimes affecting more than 60 reactions in a single individual. Importantly, these shifts did not cluster strongly by ancestry despite ancestry-associated trends in variant burden, indicating that the specific metabolic consequences of damaging variants are shaped primarily by the combination of variants within individuals rather than by continental ancestry.

The modular design of VariantFlux allows integration of omics data, enabling future exploration of gene–transcript–metabolite relationships and personalised pathway adaptation. This extensibility positions the workflow as a platform for pharmacogenomics, precision nephrology, and mechanistic interpretation of population-level genetic studies. As an example, prior work on a PAR4 variant in Tiwi Islanders^52^ highlighted the difficulty of linking single variants to kidney function without mechanistic insight. While allele frequencies differed across global populations, no association with renal function was detected—emphasising the need for tools that can reveal how specific variants influence metabolic pathways and metabolite production. VariantFlux directly addresses such gaps by mapping genetic variation to downstream metabolic consequences.

Finally, comparison of model-predicted essential genes with curated essentiality and gene–disease databases validated a substantial fraction of predictions and identified additional kidney-specific candidate genes with limited prior annotation. These genes represent promising targets for follow-up studies in renal physiology and disease, particularly where damaging variants are enriched among patient genomes.

Overall, this study reveals a dual landscape of metabolic stability and plasticity: robust preservation of global kidney metabolic capacity, coupled with widespread, ancestry-modulated, and highly individual-specific flux reorganisation. By linking human genetic diversity to mechanistic predictions of metabolic function, VariantFlux advances the resolution at which we can interpret the metabolic consequences of genomic variation and brings us closer to personalised, ancestry-aware metabolic modelling for clinical and translational applications.

---

**Supplementary Figure S1.**
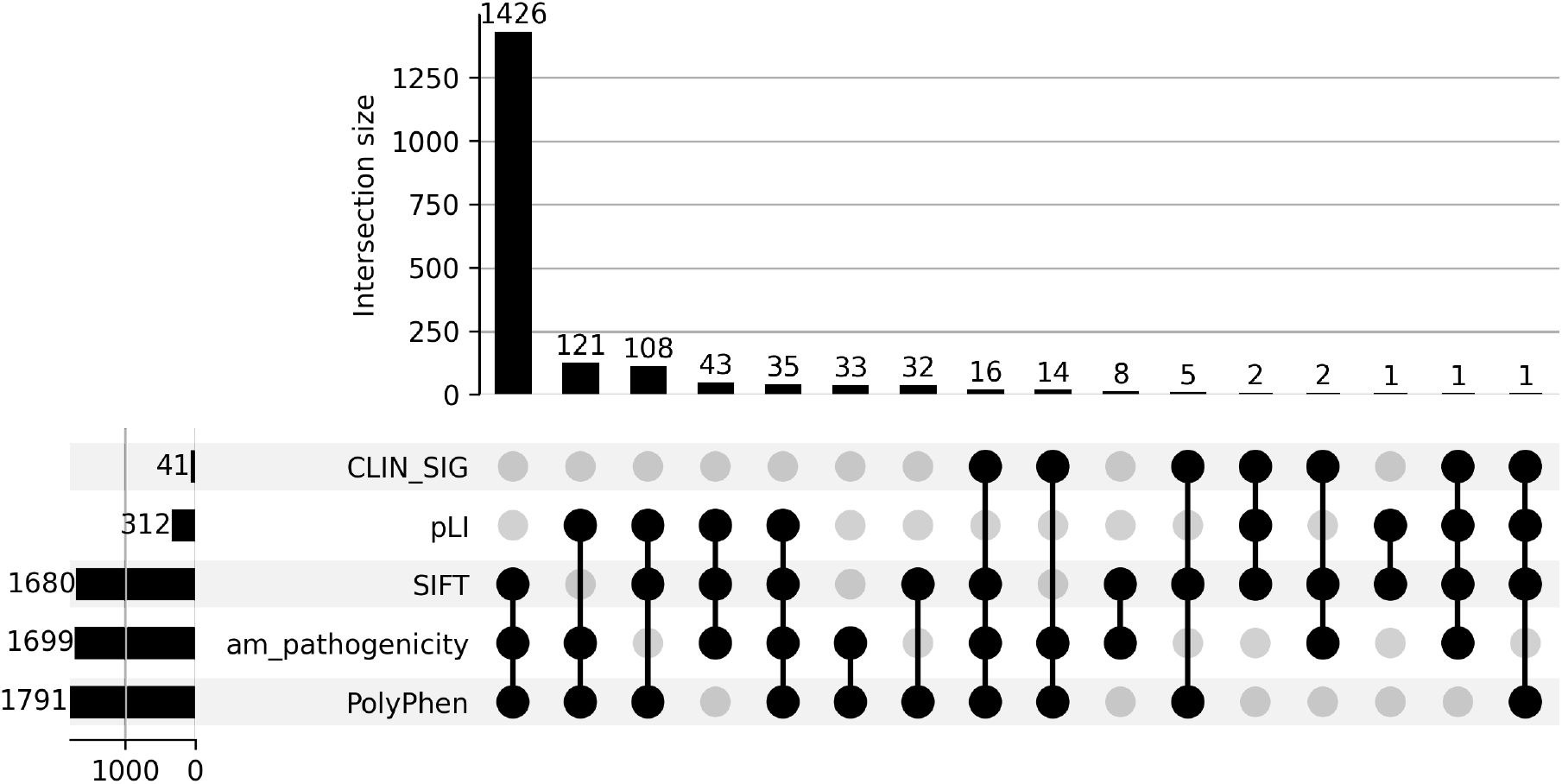
Counts of genes predicted as damaging by different variant prediction tools and for all possible combinations of prediction tools.

**Supplementary Figure S2.**
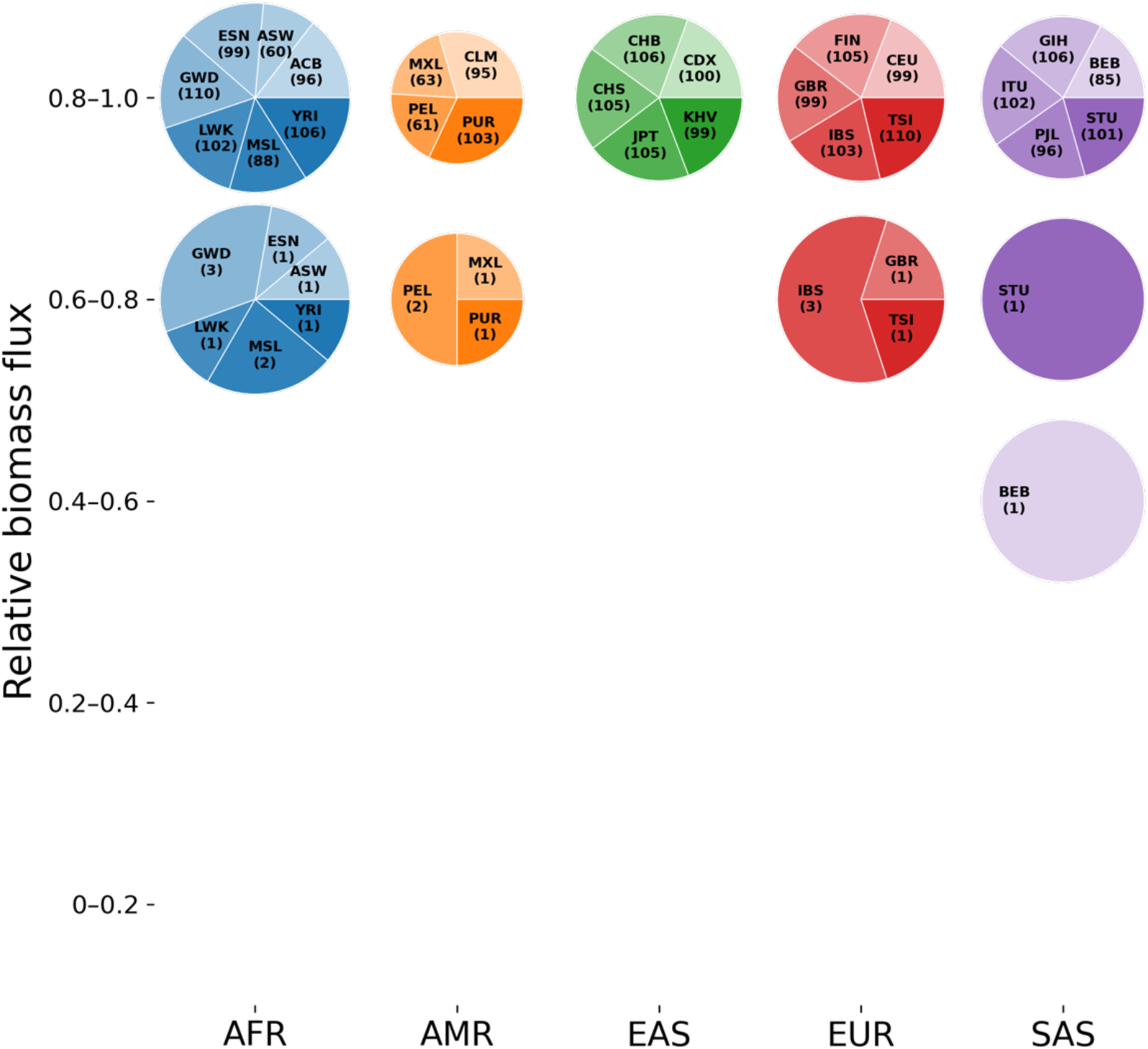
Predicted relative growth rate (i.e., biomass flux) of perturbed kidney models for 2,547 individuals across all sub-populations within each ancestry group. Relative biomass fluxes were calculated by dividing the biomass flux predicted by the perturbed model to that of the unperturbed baseline model. For visualisation purposes, relative biomass fluxes are binned into five groups of 0.2 intervals. Population abbreviations are consistent with the nomenclature used by IGSR. Numbers in parenthesis indicate the number of individuals within that population. Results are shown for simulating full knockouts of both homozygous and heterozygous states of the genes as described in Figure 3.

## References

1. Hook, P. W. & Timp, W. Beyond assembly: the increasing flexibility of single-molecule sequencing technology. Nature Reviews Genetics vol. 24 627–641 Preprint at 10.1038/s41576-023-00600-1 (2023).

2. Eveleigh, R. J. M. et al. Finding an optimal sequencing strategy to detect short and long genetic variants in a human genome. Preprint at 10.1101/2025.05.30.656631 (2025).

3. Wörheide, M. A., Krumsiek, J., Kastenmüller, G. & Arnold, M. Multi-omics integration in biomedical research – A metabolomics-centric review. Analytica Chimica Acta vol. 1141 144–162 Preprint at 10.1016/j.aca.2020.10.038 (2021).

4. Sherman, R. M. & Salzberg, S. L. Pan-genomics in the human genome era. Nature Reviews Genetics vol. 21 243–254 Preprint at 10.1038/s41576-020-0210-7 (2020).

5. Qiao, L., Khalilimeybodi, A., Linden-Santangeli, N. J. & Rangamani, P. The Evolution of Systems Biology and Systems Medicine: From Mechanistic Models to Uncertainty Quantification. 55, 26 (2025).

6. Whole-genome sequencing of 490,640 UK Biobank participants. Nature 645, 692–701 (2025).

7. Xue, Y. et al. Deleterious- and disease-allele prevalence in healthy individuals: Insights from current predictions, mutation databases, and population-scale resequencing. Am J Hum Genet 91, 1022–1032 (2012).

8. MacArthur, D. G. et al. A systematic survey of loss-of-function variants in human protein-coding genes. Science (1979) 335, 823–828 (2012).

9. Karczewski, K. J. et al. The mutational constraint spectrum quantified from variation in 141,456 humans. Nature 581, 434–443 (2020).

10. Riccio, C., Jansen, M. L., Guo, L. & Ziegler, A. Variant effect predictors: a systematic review and practical guide. Human Genetics vol. 143 625–634 Preprint at 10.1007/s00439-024-02670-5 (2024).

11. Martin, A. R. et al. Clinical use of current polygenic risk scores may exacerbate health disparities. Nat Genet 51, 584–591 (2019).

12. Reis, A. L. M. et al. The landscape of genomic structural variation in Indigenous Australians. Nature 624, 602–610 (2023).

13. Domingo-Gallego, A. et al. Clinical utility of genetic testing in early-onset kidney disease: Seven genes are the main players. Nephrology Dialysis Transplantation 37, 687–696 (2022).

14. Huerta-Chagoya, A. et al. Multi-ancestry polygenic risk scores for the prediction of type 2 diabetes and complications in diverse ancestries. Clicerio González-Villalpando 15, 185.

15. Suhre, K. & Gieger, C. Genetic variation in metabolic phenotypes: Study designs and applications. Nature Reviews Genetics vol. 13 759–769 Preprint at 10.1038/nrg3314 (2012).

16. Fountoglou, A., Deltas, C., Siomou, E. & Dounousi, E. Genome-wide association studies reconstructing chronic kidney disease. Nephrology Dialysis Transplantation vol. 39 395–402 Preprint at 10.1093/ndt/gfad209 (2024).

17. Schlosser, P. et al. Genetic studies of paired metabolomes reveal enzymatic and transport processes at the interface of plasma and urine. Nat Genet 55, 995–1008 (2023).

18. Rizi, S., Goss, N., Kutalik, Z. & van der Graaf, A. Comprehensive metabolite ratio QTL mapping reveals disease relevant enzyme biology. Preprint at 10.64898/2025.12.04.25341616 (2025).

19. Scherer, N. et al. Coupling metabolomics and exome sequencing reveals graded effects of rare damaging heterozygous variants on gene function and human traits. Nat Genet 57, 193–205 (2025).

20. Cheng, Y. et al. Rare genetic variants affecting urine metabolite levels link population variation to inborn errors of metabolism. Nat Commun 12, (2021).

21. Bordbar, A., Monk, J. M., King, Z. A. & Palsson, B. O. Constraint-based models predict metabolic and associated cellular functions. Nature Reviews Genetics vol. 15 107–120 Preprint at 10.1038/nrg3643 (2014).

22. Mardinoglu, A. & Palsson, B. Genome-scale models in human metabologenomics. Nature Reviews Genetics vol. 26 123–140 Preprint at 10.1038/s41576-024-00768-0 (2025).

23. Brunk, E. et al. Recon3D enables a three-dimensional view of gene variation in human metabolism. Nat Biotechnol 36, 272–281 (2018).

24. Lander, S., et al. Initial Sequencing and Analysis of the Human Genome International Human Genome Sequencing Consortium* The Sanger Centre: Beijing Genomics Institute/Human Genome Center. NATURE vol. 409 www.nature.com (2001).

25. Harrison, P. W. et al. Ensembl 2024. Nucleic Acids Res 52, D891–D899 (2024).

26. Danecek, P. et al. Twelve years of SAMtools and BCFtools. Gigascience 10, (2021).

27. McLaren, W. et al. The Ensembl Variant Effect Predictor. Genome Biol 17, (2016).

28. Ng, P. C. & Henikoff, S. Predicting deleterious amino acid substitutions. Genome Res 11, 863–874 (2001).

29. Adzhubei, I. A. et al. A method and server for predicting damaging missense mutations. Nature Methods vol. 7 248–249 Preprint at 10.1038/nmeth0410-248 (2010).

30. Lek, M. et al. Analysis of protein-coding genetic variation in 60,706 humans. Nature 536, 285–291 (2016).

31. Cheng, J. et al. Accurate proteome-wide missense variant effect prediction with AlphaMissense. Science (1979) 381, (2023).

32. Robinson, J. L., et al. An Atlas of Human Metabolism. Sci. Signal vol. 13 https://www.science.org (2020).

33. Wishart, D. S. et al. HMDB 5.0: The Human Metabolome Database for 2022. Nucleic Acids Res 50, D622–D631 (2022).

34. Knol, M. G. E., Wulfmeyer, V. C., Müller, R. U. & Rinschen, M. M. Amino acid metabolism in kidney health and disease. Nature Reviews Nephrology vol. 20 771–788 Preprint at 10.1038/s41581-024-00872-8 (2024).

35. Chang, R. L., Xie, L., Xie, L., Bourne, P. E. & Palsson, B. Drug off-target effects predicted using structural analysis in the context of a metabolic network model. PLoS Comput Biol 6, (2010).

36. Quek, L. E., et al. Reducing Recon 2 for steady-state flux analysis of HEK cell culture. J Biotechnol 184, 172–178 (2014).

37. KRIz, W., et al. A Standard Nomenclature for Structures of the Kidney THE RENAL COMMISSION OF THE INTERNATIONAL UNION OF PHYSIOLOGICAL SCIENCES (IUPS) Prepared by Physiology: Renal, Fluid and Electrolyte Physiology. Kidney International vol. 33 (1988).

38. Guo, C. et al. Crosstalk between proximal tubular epithelial cells and other interstitial cells in tubulointerstitial fibrosis after renal injury. Frontiers in Endocrinology vol. 14 Preprint at 10.3389/fendo.2023.1256375 (2023).

39. Edwards, J. S. & Palsson, B. O. The Escherichia Coli MG1655 in Silico Metabolic Genotype: Its Definition, Characteristics, and Capabilities. vol. 97 http://gcrg.ucsd.edudownloads.html. (2000).

40. Lance, G. N. & Williams, W. T. Computer Programs for Hierarchical Polythetic Classification (‘similarity Analyses’). http://comjnl.oxfordjournals.org/.

41. Hill-Burns, E. M. et al. Parkinson’s disease and Parkinson’s disease medications have distinct signatures of the gut microbiome. Movement Disorders 32, 739–749 (2017).

42. Levy, A., Shalom, B. R. & Chalamish, M. A guide to similarity measures and their data science applications. J Big Data 12, (2025).

43. Lewis, N. E. et al. Omic data from evolved E. coli are consistent with computed optimal growth from genome-scale models. Mol Syst Biol 6, (2010).

44. Orth, J. D., Thiele, I. & Palsson, B. O. What is flux balance analysis? Nature Biotechnology vol. 28 245–248 Preprint at 10.1038/nbt.1614 (2010).

45. Ebrahim, A., Lerman, J. A., Palsson, B. O. & Hyduke, D. R. COBRApy: COnstraints-Based Reconstruction and Analysis for Python. BMC Syst Biol 7, (2013).

46. Gurumayum, S. et al. OGEE v3: Online GEne Essentiality database with increased coverage of organisms and human cell lines. Nucleic Acids Res 49, D998–D1003 (2021).

47. DiStefano, M. T. et al. The Gene Curation Coalition: A global effort to harmonize gene–disease evidence resources. Genetics in Medicine 24, 1732–1742 (2022).

48. Martin, A. R. et al. PanelApp crowdsources expert knowledge to establish consensus diagnostic gene panels. Nature Genetics vol. 51 1560–1565 Preprint at 10.1038/s41588-019-0528-2 (2019).

49. Gargano, M. A. et al. The Human Phenotype Ontology in 2024: phenotypes around the world. Nucleic Acids Res 52, D1333–D1346 (2024).

50. Altshuler, D. M. et al. An integrated map of genetic variation from 1,092 human genomes. Nature 491, 56–65 (2012).

51. Sudmant, P. H. et al. An integrated map of structural variation in 2,504 human genomes. Nature 526, 75–81 (2015).

52. Ningtyas, D. et al. Analysis of the F2LR3 (PAR4) Single Nucleotide Polymorphism (rs773902) in an Indigenous Australian Population. Front Genet 11, (2020).

